# Estimated limits of organism-specific training for epitope prediction

**DOI:** 10.1101/2021.11.02.466801

**Authors:** Jodie Ashford, Felipe Campelo

## Abstract

**Background:** The identification of linear B-cell epitopes remains an important task in the development of vaccines, therapeutic antibodies and several diagnostic tests. Machine learning predictors are trained to flag potential epitope candidates for experimental validation and currently, most predictors are trained as generalist models using large, heterogeneous data sets. Recently, organism-specific training has been shown to improve prediction performance for data-rich organisms. Unfortunately, for most organisms, large volumes of validated epitope data are not yet available. This article investigates the limits of organism-specific training for epitope prediction. It explores the validity of organism-specific training for data-poor organisms by examining how the size of the training data set affects prediction performance. It also compares the performance of organism-specific training under simulated data-poor conditions to that of models trained using traditional large heterogeneous and hybrid data sets.

**Results:** This work shows how models trained on small organism-specific data sets can outperform similar models trained on (potentially much larger) heterogeneous and mixed data sets. The results reported indicate that as few as 20 labelled peptides from a given pathogen can be sufficient to generate models that outperform widely-used predictors from the literature, which are trained on heterogeneous data. Models trained using more than about 100 to 150 organism-specific peptides perform consistently better than most generalist models across a wide variety of performance measures, and in some cases can even approach the performance of organism-specific models trained on considerably larger data sets.

**Conclusions:** Organism-specific training improves linear B-cell epitope prediction performance even in situations when only small training sets are available, which opens new possibilities for the development of bespoke, high-performance predictive models when studying data-poor organisms such as emerging or neglected pathogens.

## Background

The immune system is a complex network of processes designed to protect the body against pathogens. One vital aspect of human immunity is humoral, or antibody-mediated, immunity. In humoral immunity, B-lymphocytes, also known as B-cells, are activated when their B-cell receptor (BCR) binds with an antigen. Activated B-cells then produce antibodies which are released into the circulatory system to find and bind with their specific antigens [1]. This antigen-antibody recognition is a vital process in protecting the body against pathogens and B-cells are key cells in this process.

A B-cell epitope (or antigenic determinant) is the exact portion of an antigen that the antigen-binding site of a B-cell receptor recognises and binds to [2, 3]. B-cell epitope identification is an essential process in a number of medical processes; it can help with therapeutic antibody production, vaccine development and in developing diagnostic tools [4–6]. There are two categories of epitopes: linear and conformational. Linear or continuous epitopes correspond to contiguous sequences of amino acid (AA) residues; these epitopes are recognised by antibodies by their primary structure/linear sequence of amino acids. Conformational or discontinuous epitopes are formed by AAs that, although separated in the primary sequence, are brought together by protein folding [7, Chapter 3].

There are several different types of epitope prediction methods and the type of method deployed may differ depending on the type of epitope (linear or conformational) being predicted. Most current epitope prediction methods are designed to predict linear epitopes [8–18], though the majority of epitopes (∼90% of all B-cell epitopes) are thought to be conformational [19, 20]. There are multiple reasons for this: due to their nature linear epitopes can be predicted from protein sequence data alone, which is readily available in numerous public databases [21–24]; conformational epitopes, on the other hand, require structural protein data for prediction which is not as readily available. Predicting conformational epitopes also takes more time as it is more computationally expensive than linear epitope prediction [25] and these epitopes are more difficult to synthesise in the laboratory [26, Chapter 1]. For these reasons many epitope prediction works, including ours, focus on linear B-cell epitope prediction.

Traditionally, experimental methods were used for B-cell epitope identification, for example: X-ray crystallography, peptide arrays, enzyme-linked immunosorbent assay (ELISA) and phage display [26–28]. However, these methods are time consuming, resource intensive and technically difficult to execute [6,26]. Because of this and the current availability of protein sequence data the focus is now on computational methods for epitope prediction. Machine learning algorithms for epitope prediction are trained to be able to distinguish B-cell epitopes from non-epitopes. Numerous ML methods exist for B-cell epitope prediction and these methods have been shown to outperform early epitope prediction methods based solely on simple amino acid propensity scale calculations, though this is not always the case [3, 29]. Examples of machine learning approaches for epitope prediction include: neural network-based methods such as ABCpred [12], which uses a recurrent neural network (RNN) to predict B-cell epitopes from antigen sequences using fixed length patterns and other amino acid composition-based features as input. Other popular ML methods for epitope prediction include Support Vector Machines [30] which have been used in many epitope prediction pipelines [13, 31–39]. One example of this is BCPred [13], which uses Support Vector Machine (SVM) classifiers with string kernels [13]. Random Forest Classifiers [40] have also been used in multiple epitope prediction pipelines [17, 41], including the work by Saravanan and Gautham [42] which described an amino acid composition-based feature descriptor, Dipeptide Deviation from Expected Mean (DDE), and evaluated it using a support vector machine and an AdaBoost-Random Forest, with the latter exhibiting the best performance.

ML methods like the ones mentioned above help to bypass some of the difficulties usually encountered by traditional experimental epitope prediction methods [25,43,44]. However, there is room for improvement as many prediction methods still exhibit relatively low prediction performance [25]. Currently, most epitope prediction models are trained on large heterogeneous data sets made up of observations from multiple organisms including: prokaryotes, viruses, fungi, protozoan, humans and other eukaryotes. However, in a recent work [45] we have shown that training models on smaller organism-specific data sets may improve prediction performance. In that work, organism-specific models were developed for three different organisms: the nematode *Onchocerca volvulus*, the Epstein-Barr Virus and the Hepatitis C Virus. These organisms were selected due to the availability of a large volume of observations (both validated positive and negative epitopes) for them in the Immune Epitope Database (IEDB) [46]. The results obtained showed that, for these data-rich organisms, organism-specific models outperformed models trained on much larger heterogeneous data sets as well as several of the best epitope prediction tools from the literature, across multiple performance measures.

Unfortunately, large volumes of validated epitope data are not available for most organisms - either because the organism relates to a neglected or emerging disease, or because it may have only a small number of epitopes. The aim of this study is to investigate the limits of organism-specific training, by focusing on two main questions: (i) How does the number of available organism-specific training peptides affect prediction performance?; and (ii) What is the smallest volume of organism-specific data that produces models surpassing the performance of those trained on large, heterogeneous data sets? To answer those questions, we calculate and compare the prediction performance of models trained on reduced training sets against models trained on mixed data as well as on large, heterogeneous data sets. We also contrast the observed performances with four predictors from the literature trained as generalist (as opposed to organism-specific) models – Bepipred2.0 [17], LBtope [39], iBCE-EL [47] and ABCpred [6] – across multiple performance measures.

## Methods

### Data Sets

Data from three pathogens was taken from [45], specific to the organisms: *Onchocerca volvulus* (taxonomy ID: 6282;), Epstein-Barr Virus (taxonomy ID: 10376) and Hepatitis C Virus (taxonomy ID: 11102). These data sets were generated based on the full XML export of the IEDB retrieved on the 10th of October 2020, and filtered according to the criteria listed in [45] (section 2.1, “Data sets”). The available data was split at the protein level, with entries coming from the same protein, or from proteins exhibiting sequence coverage and similarity greater than 80%, always placed in the same split. Two base sets were derived from the data available for each organism: a *Hold-out* set containing approximately 25% of the data; and a second set containing the remaining observations to be used for all model development activities. A set of *Heterogeneous* data was also extracted for each organism, by randomly sampling observations, grouped by taxonomy ID, from the full IEDB export (excluding any observations related to the specific organism). These heterogeneous sets contain around 6000 labeled peptides, with a 50% class balance.

To investigate the effect of the size of organism-specific data sets on prediction performance, and try to estimate rough lower bounds of the required amount of data for organism-specific training to still represent a good alternative to models developed on larger, heterogeneous data sets, we extracted several *reduced* organism-specific and heterogeneous/hybrid training sets for each organism, based on the available model development data described above. For each organism and each desired training set size, we split the full model development data into smaller non-overlapping *Organism-specific* data sets, each containing data from between 20 and 500 peptides (see figure 1). The same class balance as the full organism-specific data set was maintained in all subsets.

**Figure 1:**
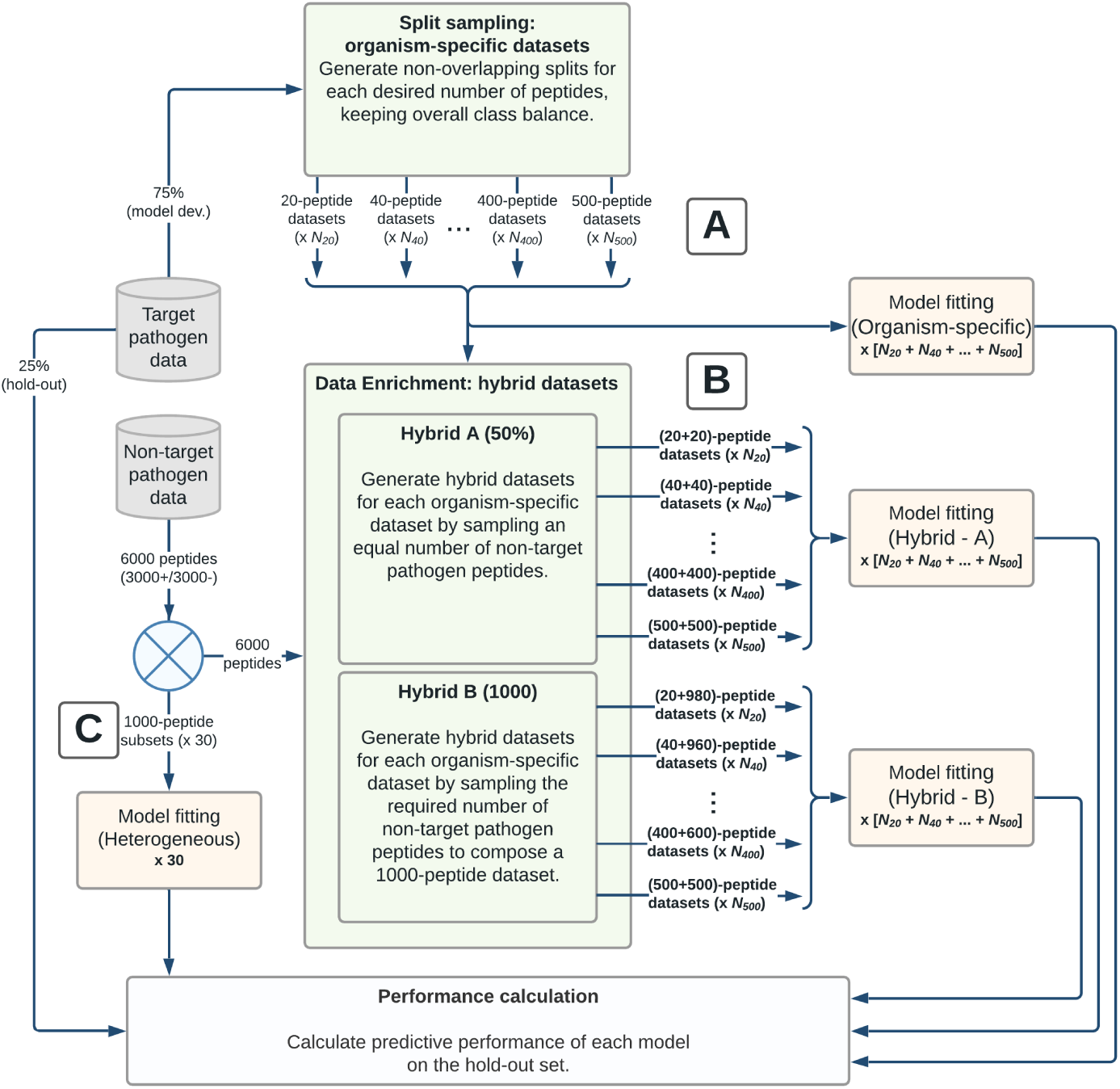
Experimental protocol for testing the limits of organism-specific model training for linear B-cell epitope prediction. (A) For each pathogen and each desired data size (in terms of number of peptides from the target pathogen), the model development data is split into non-overlapping subsets of the desired size, each maintaining the original class balance of the data. (B) Two sets of hybrid data sets are composed based on the organism-specific reduced-data replicates: *Hybrid-A* maintains a fixed 50-50 balance between organism-specific and heterogeneous data at all data set sizes; *Hybrid-B* adds the required number of non-target organism observations to complete a data set of 1,000 peptides, and therefore results in sets with variable proportions of organism-specific peptides. (C) Baseline data sets composed of 1,000 exclusively non-target pathogen peptides are also generated based on different sub-samplings (without replacement) from the heterogeneous data. All data sets are used to train Random Forest models, which then have their performances assessed on organism-specific hold-out data.

Table 1 details the information on the reduced organism-specific data sets generated for each pathogen. Based on these variable-sized organism-specific training sets, we assembled two groups of *hybrid* data sets:

**Table 1:** Summary information on the organism-specific data sets of each pathogen: number of positive/negative peptides in the hold-out and model development sets, and number of replicates for each set size (set sizes being defined by the number of organism-specific peptides in the set). Hybrid-A and Hybrid-B sets were generated based on the same subsets of organism-specific peptides, and therefore have the same number of replicates at each size. Heterogeneous sets were generated separately, with 30 replicates of 1,000 non-target pathogen peptides used in the experiments.

|  | *O. volvulus* | Hepatitis C virus | Epstein-Barr virus |
| --- | --- | --- | --- |
| Hold-out peptides | (832+ / 777-) | (218+ / 358-) | (625+ / 315-) |
| Model dev. peptides | (2441+ / 2378-) | (919+ / 783-) | (1746+ / 811-) |
| 20-peptide sets ($N_{20}$) | 237 | 83 | 124 |
| 40-peptide sets ($N_{40}$) | 118 | 41 | 62 |
| 60-peptide sets ($N_{60}$) | 79 | 27 | 42 |
| 80-peptide sets ($N_{80}$) | 59 | 21 | 31 |
| 100-peptide sets ($N_{100}$) | 47 | 17 | 25 |
| 150-peptide sets ($N_{150}$) | 32 | 11 | 16 |
| 200-peptide sets ($N_{200}$) | 24 | 8 | 12 |
| 250-peptide sets ($N_{250}$) | 19 | 6 | 10 |
| 300-peptide sets ($N_{300}$) | 16 | 5 | 8 |
| 400-peptide sets ($N_{400}$) | 12 | 4 | 6 |
| 500-peptide sets ($N_{500}$) | 9 | 4 | 5 |

**Table 2:**
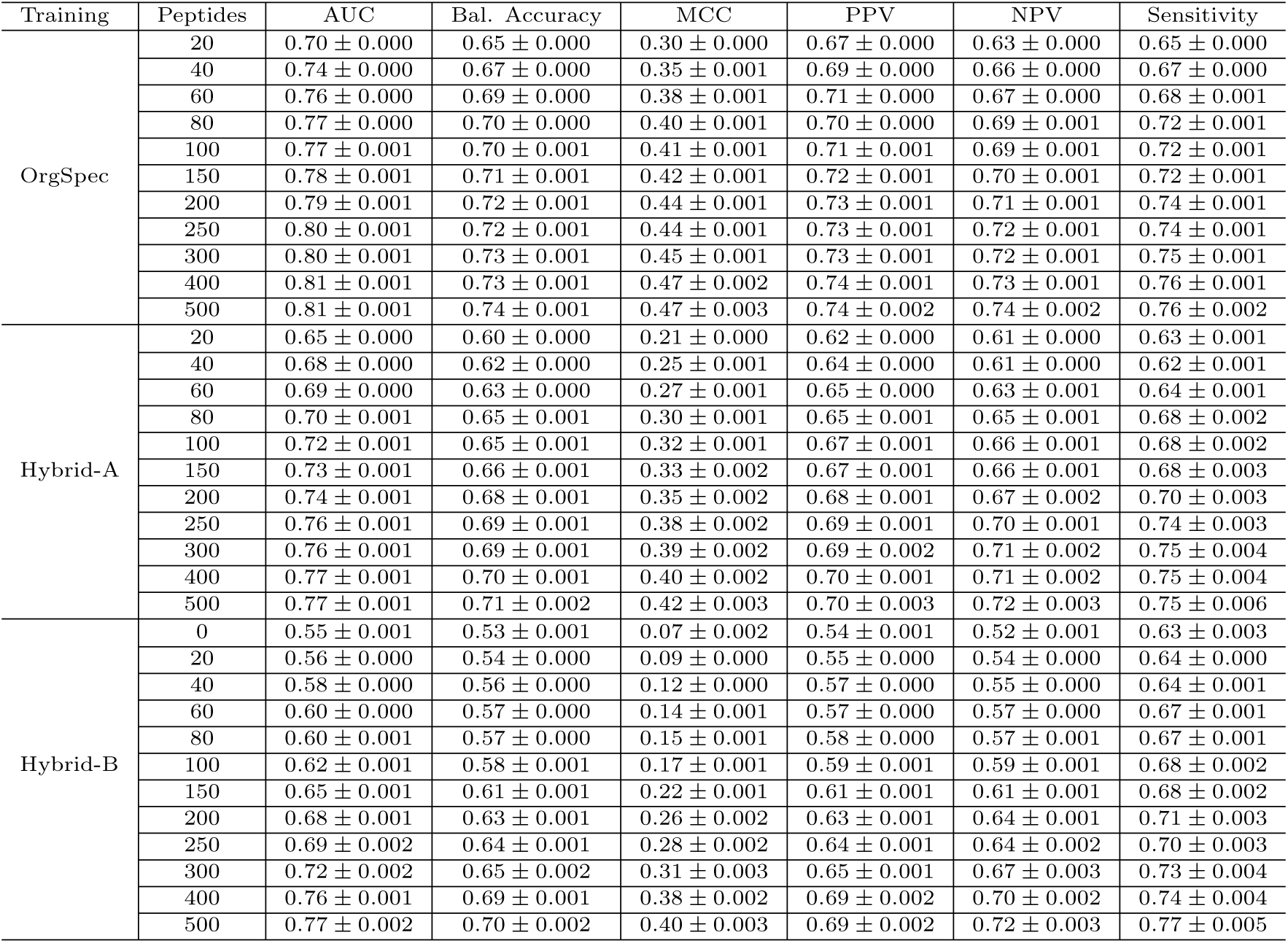
Point estimates and standard errors of mean performance for models trained on the *Onchocerca volvulus* data. Row *Hybrid-B* : 0 Peptides corresponds to the “heterogeneous” case, with 0 organismspecific and 1,000 non-target pathogen peptides in the training set.

**Table 3:**
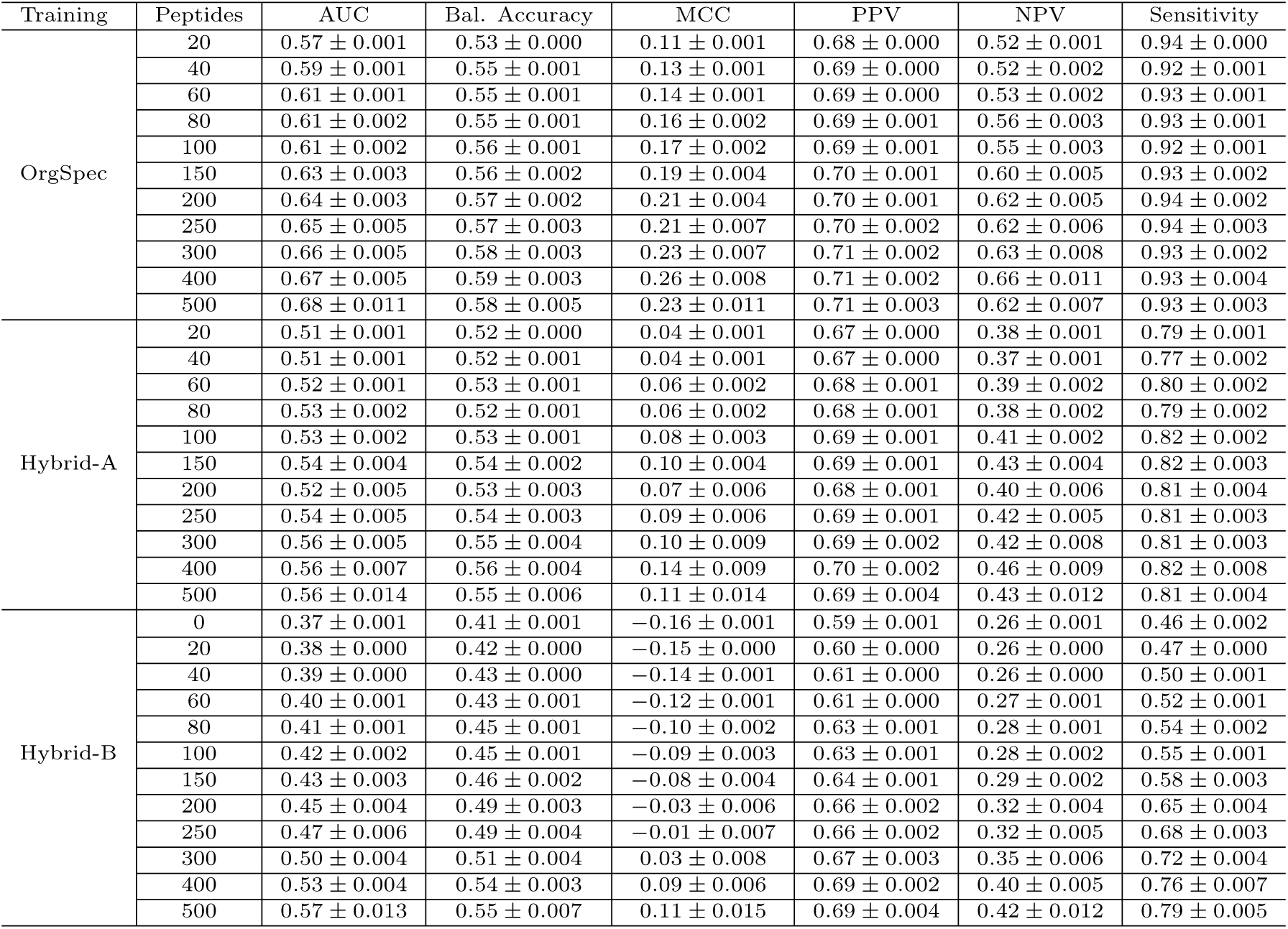
Point estimates and standard errors of mean performance for models trained on the Epstein-Barr Virus data. Row *Hybrid-B* : 0 Peptides corresponds to the “heterogeneous” case, with 0 organism-specific and 1,000 non-target pathogen peptides in the training set.

**Table 4:**
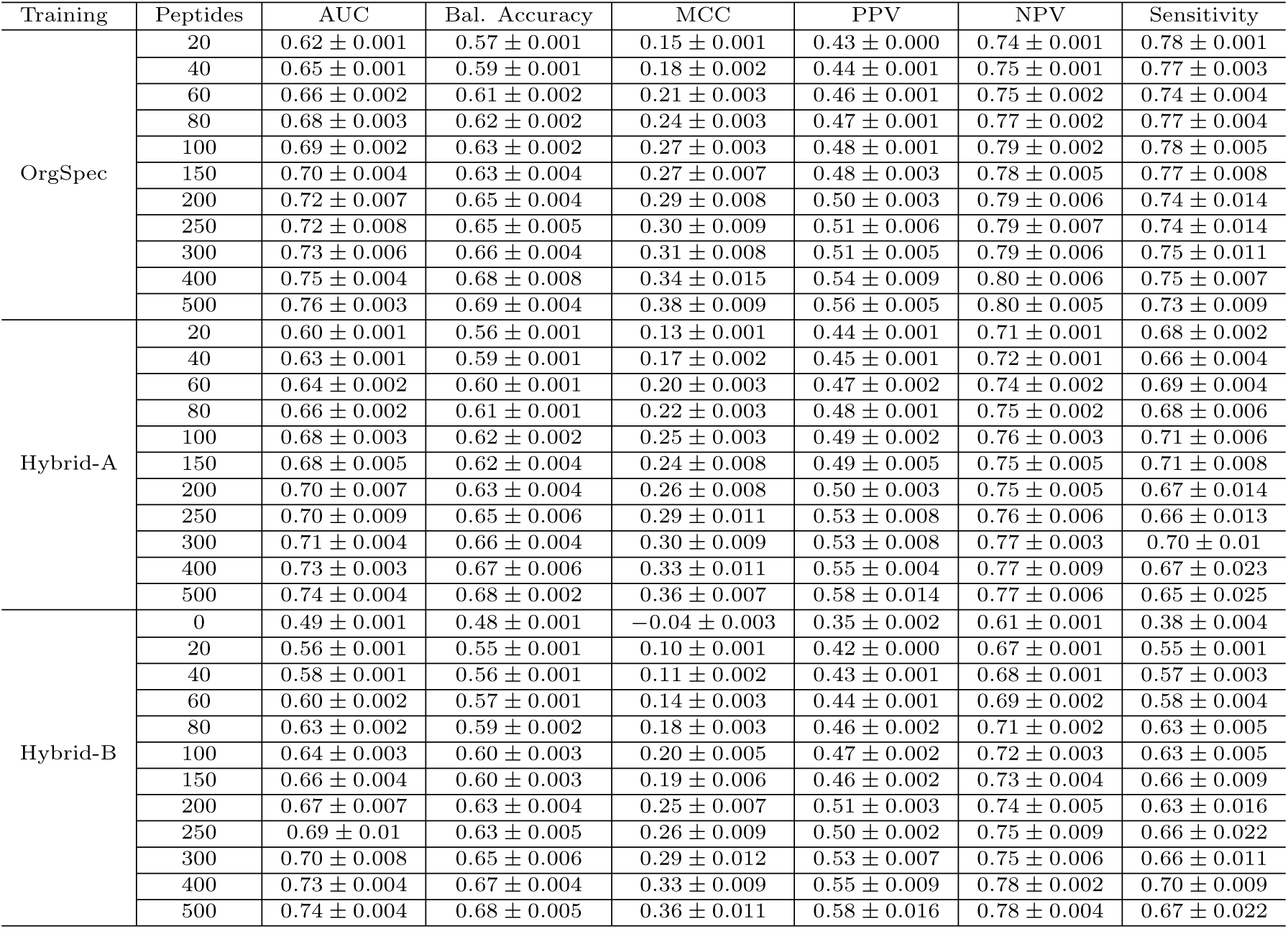
Point estimates and standard errors of mean performance for models trained on the Hepatitis C Virus data. Row *Hybrid-B* : 0 Peptides corresponds to the “heterogeneous” case, with 0 organism-specific and 1,000 non-target pathogen peptides in the training set.

- *Hybrid-A*, composed of the organism-specific peptides plus an equal amount of peptides sampled from other pathogens. Consequently, *Hybrid-A* data sets were always composed of twice as many peptides as their corresponding organism-specific ones, and the balance between organism-specific and “other” peptides was always 50-50%.
- *Hybrid-B*, composed of the organism-specific peptides plus the required amount of peptides sampled from other pathogens to complete a data set size of 1, 000 training peptides (e.g., 20 organism specific + 980 “other” peptides, 40+960, etc.). *Hybrid-B* data sets had a fixed size, but a varying balance of data from the target pathogen vs. other organisms.

For each data size tested (defined in this experiment as the number of organism-specific peptides in the data sets) both the *Hybrid-A* and *Hybrid-B* groups had the same number of replicates as the organism-specific sets of that size. The number of replicates at each size is documented in Table 1.

Besides the hybrid data sets, for each target pathogen we also fit models on 30 samples of 1, 000 peptides from “other” organisms. In the results this is analysed as the limit case of the *Hybrid-B* data sets (as a “0+1000” -peptide set). Figure 1 illustrates the full experimental pipeline, including the generation of all relevant data sets.

### Modelling and Performance Assessment

Epitope prediction models were developed by training Random Forest (RF) predictors on each of the training data sets outlined above, using Scikit-learn version 0.24.1 [48] under standard hyper-parameter values. The choice of Random Forest was based on preliminary experimentation, as documented in [45], and also to make this work more directly comparable with the results reported in that earlier one. The trained models were then used to generate predictions for the organism-specific hold-out data sets and prediction performance was assessed using multiple different performance measures, namely: Balanced Accuracy (BAL.ACC), Matthew’s Correlation Coefficient (MCC), Area Under the Curve (AUC), Positive Predictive Value (PPV), Negative Predictive Value (NPV) and Sensitivity (SENS). Being as these measures were calculated on the hold-out data sets (which were not seen by the models at any point other than testing) it can be assumed that these values represent a reasonable estimate of the generalisation performance of the models used for epitope prediction on proteins coming from each of the pathogens. The estimated mean performance and standard errors for each quality indicator were calculated from the replicates at each pathogen and data set size.

Besides the new results obtained by the models trained in this study, we also included the following comparison baselines, extracted from [45], in our assessment of the models’ performance: the observed performance of Bepipred2.0 [17], LBtope [39], iBCE-EL [47] and ABCpred [6], on the hold-out set of each pathogen; the results obtained by Random Forest models trained on the full model development data and on a set of 6000 non-target pathogen peptides, for each organism.

## Results

Figures 2-3 display the mean performance results from each set of models on the hold-out data set of each pathogen. Each figure plots the number of organism-specific peptides in the training data set *versus* the estimated mean performance according to different indicators.

**Figure 2:**
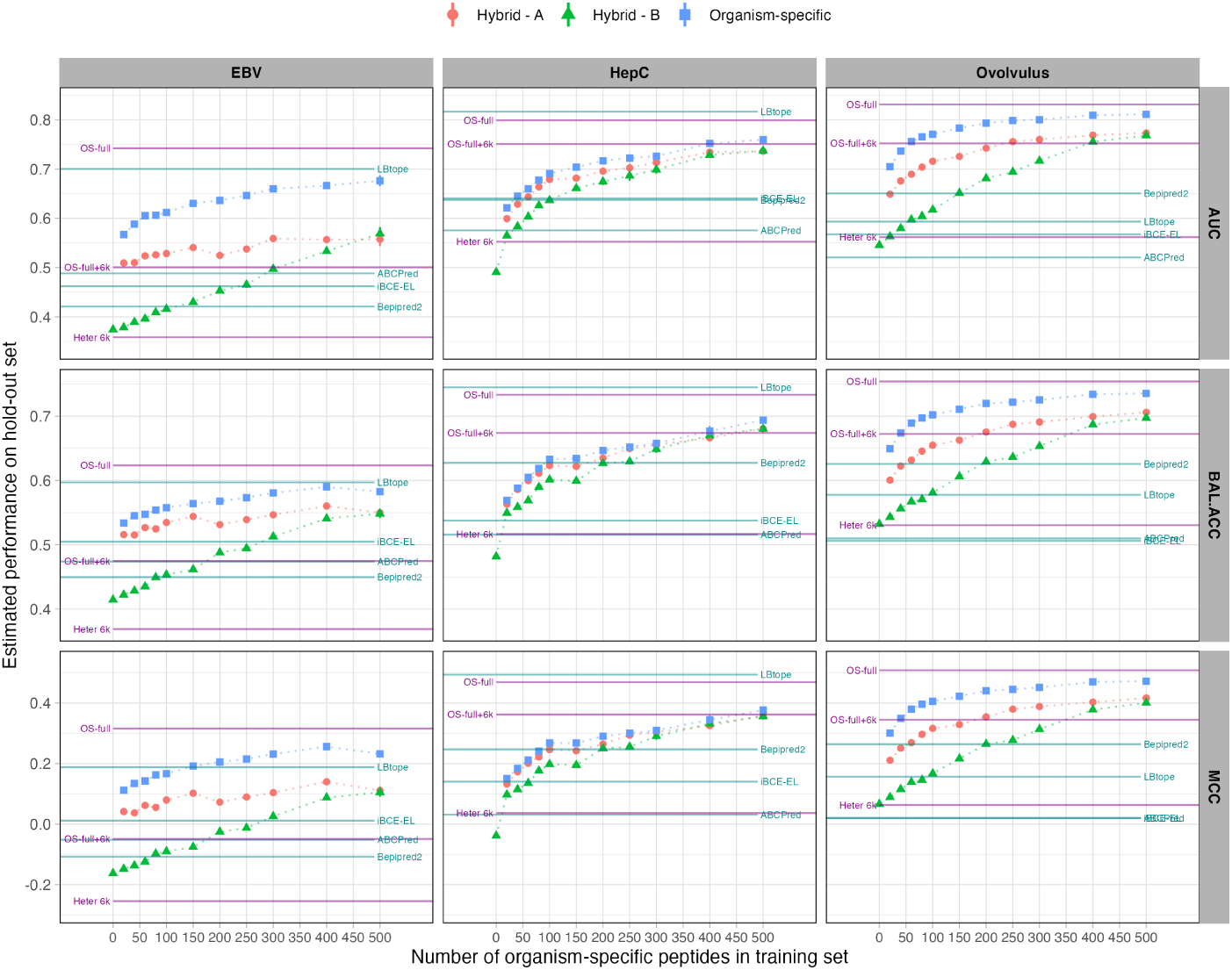
Mean performance (AUC, balanced accuracy, MCC) and standard errors for all models tested. Blue squares indicate the scores from models trained on organism-specific data sets, red circles are from those trained on the *Hybrid-A* (“doubled data”) data sets, and green triangles refer to models trained on the *Hybrid-B* (“1000 peptides”) data sets. Horizontal lines indicate reference values extracted from [45]: models trained on the full training set (*‘OS-full’*), on a large heterogeneous set (*‘Heter 6K’*), and on a large hybrid set (*‘OSfull+6K’*), as well as the scores of several predictors from the literature on the same hold-out sets. For all pathogens tested, organism-specific training resulted in uniformly better performance across all data sizes when compared to models trained on hybrid or purely heterogeneous data, even when as few as 20 organism-specific peptides are used in the training set. Notice also how the performance of organism-specific models quickly surpasses that of most of the comparison predictors tested, even when very few organism-specific peptides are available to fit the models. (Note: standard error bars are in most cases shorter than the size of the point estimate markers)

**Figure 3:**
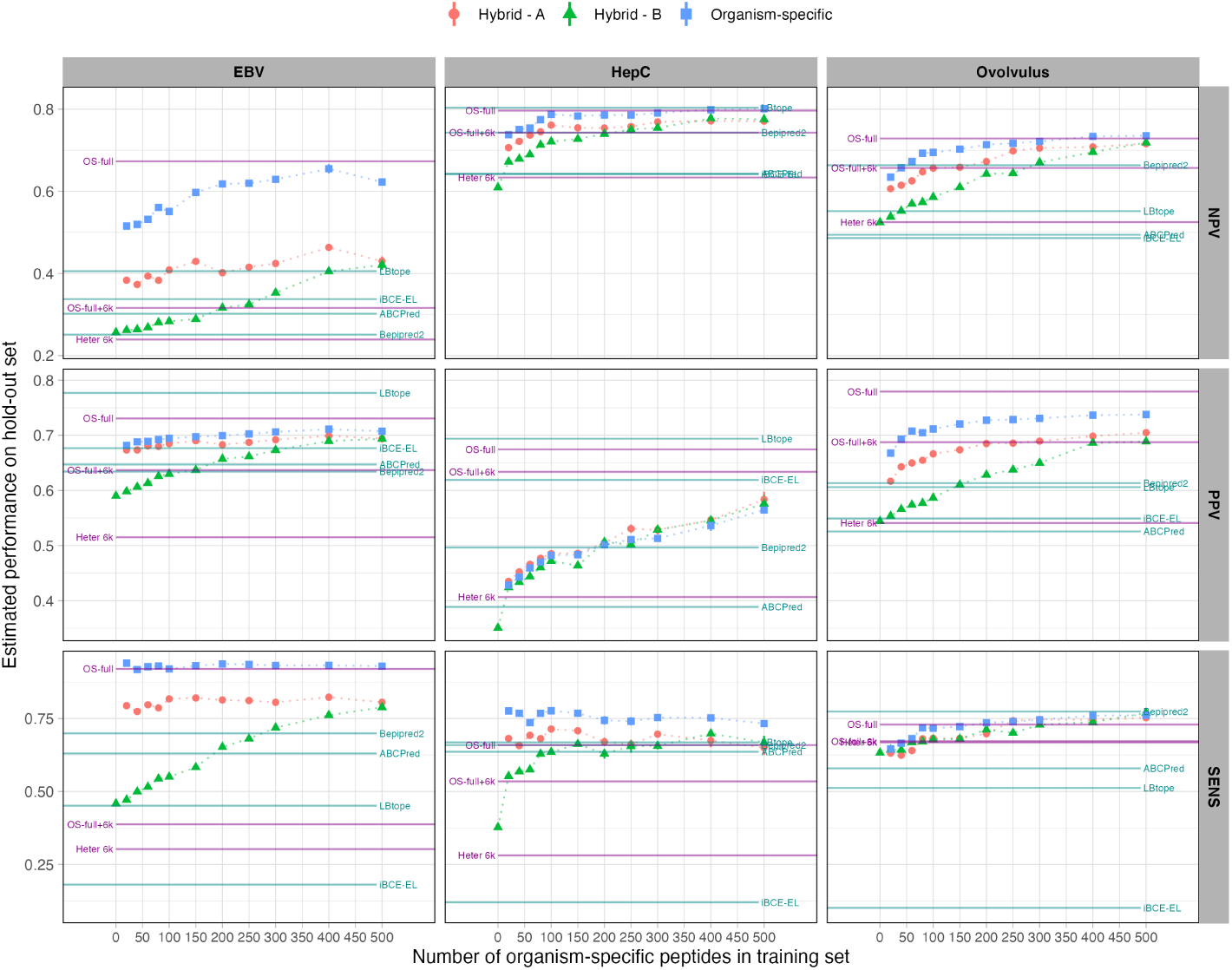
(Continuing from figure 2) Mean performance (PPV, NPV, Sensitivity) and standard errors for all models tested. The same pattern observed in figure 2 – uniform superiority of organism-specific training, when compared to models trained on bigger, but hybrid/heterogeneous, data sets – is also observed in the case of the three performance metrics shown here. As documented in [45], the apparently excellent performance of LBtope on the Hepatitis C data across all performance indicators can be partially attributed to the fact that several of the hold-out peptides used in this work are also found in LBtope’s training data [39].

For *Onchocerca volvulus* (the largest data set in this study), the highest scores on the hold-out set are from the full organism-specific model (except for sensitivity), as documented earlier in [45]. The next highest scores are from the split-sampling organism-specific models (for data sizes ≥ 40 peptides), which approach the full organism-specific performance after about 150 peptides. A clear pattern can be seen across all performance measures: models trained on the organism-specific data sets consistently and uniformly outperform those trained on *Hybrid-A* (double size) data sets, which in turn outperform the models trained on *Hybrid-B* (1,000-peptides) data sets.^1^ Within each group of models tested (trained on organism-specific, *Hybrid-A* and *Hybrid-B*) the pattern of performance improvement as the training set becomes larger was observed, as expected. The small-sample organism-specific models outperform those trained on the large heterogeneous (*Heter 6k*) and large hybrid (*OS-full+6k*) models (for≥ 40 peptides), and also all models from the literature across all performance measures – except sensitivity, where Bepipred2.0 had the highest score; and NPV, where Bepipred2.0 outperformed the models trained with ≤ 40 organism-specific peptides.

A similar pattern can be noticed for the Epstein-Barr Virus data. Figure(s) 2-3 again show that, across all performance measures, the highest scores on the hold-out set are from the full organism-specific model, with the exceptions of positive predictive value (where LBtope has the highest score) and sensitivity, where the full organism-specific model and the reduced split-sampling organism-specific models have very similar scores across all training data sizes. The second highest scores across all performance measures are almost always the split-sampling organism-specific models (down to the smallest size: 20 peptides) apart from for AUC, where LBtope approaches the performance of the full organism-specific model. The overall pattern of our reduced data set EBV models is the same as that of the *Onchocerca volvulus* models: organism-specific *> Hybrid-A > Hybrid-B*. The performance of the models also generally decreases as the training data becomes more scarce, as expected. The EBV small-sample organism-specific models outperform all the tested models from the literature, as well as the *OS-full+6k* & *Heter 6k* results, across all performance measures except AUC and PPV, where LBtope yields better performane values.

The results for the Hepatitis C Virus reinforce the performance patterns observed for the other two pathogens. As documented in [45], the apparently excellent performance of LBtope for this pathogen across all performance indicators can be partially attributed to the fact that several of the hold-out peptides used in this work are also part of LBtope’s training data^2^. With the exception of LBtope’s results, the pattern we observe for the Hepatitis C models closely mirrors the results on the other two pathogens, with organism-specific models generally outperforming the literature predictors tested even when trained with a very modest amount of peptides between 40 and 100, depending on the performance indicator. For this pathogen, the performance difference between each group is considerably smaller than the differences that can be seen for the other organisms tested, albeit still with a clear trend of organism-specific *> Hybrid-A > Hybrid-B* for all performance indicators except PPV, in which the three training regimens generally overlap across all data sizes.

When comparing all organism-specific reduced-data models scores to the heterogeneous model scores, across all organisms and for all performance measures, Figures 2-3 clearly show that almost all organism-specific models score considerably higher than the purely heterogeneous models (left-most point in the *Hybrid-B* group), as well as the hybrid models (*Hybrid-A* & *Hybrid-B*); the larger hybrid model from the previous study (*OS-full+6k*) and the generalist predictors from the literature, even when the organism-specific models are trained on modest-sized data sets. In all cases, prediction performance decreases as the number of organism-specific peptides used is reduced, even if the total number of peptides in the training set is kept fixed (*Hybrid-B*). For the organisms in this study, the organism-specific models also appear to be the most robust, with smaller performance decreases as the amount of organism-specific data is reduced when compared to *Hybrid-A* and, in particular, *Hybrid-B*.

## Discussion

The results from this study indicate that, when compared to heterogeneous and hybrid training, organism-specific training produces higher linear B-cell epitope prediction performance scores, *even for very small data set sizes*. The number of organism-specific peptides in the training set is shown to strongly affect the predictive performance of organism-specific models across multiple performance indicators, particularly up to 100 or 150 peptides, after which performance continues to increase with more data but with diminishing returns, asymptotically approaching that of models trained on the full available training data for each pathogen [45]. The results also show that organism-specific training outperforms generalist training (predictors from the literature, trained on peptides from a wide variety of pathogens) even when very small organism-specific data sets are available. The only systematic exception was the high observed performance of LBtope for the Hepatitis C Virus; However, as mentioned earlier, “*part of the hold-out examples used to asses the performance of the models is present in the training data of LBtope (9*.*59% of the Hep C hold-out sequences are present in the LBtope training data set)*” [45], which in the case of our experiments would result in some level of data leakage [49]. In addition to showing that organism-specific training outperforms heterogeneous and hybrid training, this work shows that adding unrelated data to organism-specific training sets *decreases* the generalisation performance of the resulting model when tasked with predicting epitopes for the target pathogen. It is also apparent that the more heterogeneous data is added to the training set, the poorer the prediction performance becomes, which can be clearly seen from the comparison between *Hybrid-A* and *Hybrid-B* results in Figures 2-3. This suggests that, when training models for organism-specific predictions, the training data sets should be as specific (containing only labelled peptides from that organism) as possible.

Taken together, the results presented here provide a strong indication that organism-specific models trained on data sets beyond 100-150 peptides provide very competitive predictive performance when compared to the generalist predictors tested. Additionally, the point at which organism-specific models start to outperform generalist predictors depends on the organism. For *O. volvulus* and Epstein-Barr Virus models the performance of organism-specific models compared favourably to that of generalist models down to the smallest organism-specific data set tested (20 peptides), while for Hepatitis C more peptides were required for the organism-specific training to become competitive. This highlights the strengths of organism-specific training and extends the conclusions and scope of application of the methods described in our previous study [45], which were limited to data-rich organisms. In contrast, this study has shown that organism-specific training improves epitope prediction performance for data-poor organisms as well. As a comparison, the number of labelled peptide examples in the full training sets used in [45] were: 8,819 for *O. volvulus*, 2,557 for Epstein-Barr Virus, and 1,702 for Hepatitis C Virus. These are three of the most data-rich organisms on the IEDB. Currently, most organisms have far fewer labelled epitope examples available to them, and this work has shown that, for many if not most of these organisms, organism-specific training can provide significant improvements in prediction performance.

## Conclusions

In a previous work, we showed that organism-specific training improves linear B-cell epitope prediction performance for data-abundant organisms. This work extends the scope of organism-specific modelling by showing that, contrary to our initial assumptions, organism-specific training is also a viable option for relatively data-poor organisms. However, it is clear that there are limits to organism-specific training for epitope prediction. The results documented in this study suggest that organism-specific models trained with more than about 100 labelled peptides will generally compare favourably to generalist predictors trained on substantially larger, but heterogeneous, data sets. It also confirms that predictive performance, across a wide variety of indicators, tends to increase monotonically with the number of organism-specific peptides included in the training data. It should be noted, however, that the results documented in this work have only been validated for reasonably class-balanced data sets. We have not tested models trained on strongly imbalanced data (the worst case among the pathogens tested was the Epstein-Barr virus data with a 2:1 balance of classes, which does not configure extreme class imbalance). While a further investigation of imbalanced classification approaches for epitope prediction would potentially help extend the scope of the organism-specific training framework even further, the results presented here, coupled with the increasingly cheap availability of computing power, already indicate a promising new direction for the development of bespoke predictors for pathogens under study, even for relatively data-poor organisms such as neglected pathogens or emerging health threats.

## Author’s contributions

JA implemented the Random Forest models and the experimental code, and contributed to the analysis and discussion of results. FC designed the experiment and implemented the analysis and visualisation routines. Both authors contributed equally to manuscript writing and review.

## Acknowledgements

We would like to thank our collaborators Dr. Francisco Lobo (UFMG, Brazil) and Dr. João Reis-Cunha (University of York, UK) for excellent discussions and insights that contributed to the ideas explored in this work.

## Footnotes

1 The largest data sets from both *Hybrid-A* and *Hybrid-B* always have very similar scores, as expected as in both cases the sets contain 500 peptides coming from the target pathogen data and 500 coming from the non-target pathogen data.

2 https://webs.iiitd.edu.in/raghava/lbtope/data/LBtope_Variable_Positive_epitopes.txt

